# Effects of warming winter embryo incubation temperatures on larval cisco (*Coregonus artedi*) survival, growth, and critical thermal maximum

**DOI:** 10.1101/2021.07.01.450800

**Authors:** Taylor R. Stewart, Mark R. Vinson, Jason D. Stockwell

## Abstract

Freshwater whitefishes, Salmonidae Coregoninae, are cold stenothermic fishes of ecological and socio-economic importance in northern hemisphere lakes that are warming in response to climate change. To address the effect of warming waters on coregonine reproduction we experimentally evaluated different embryo incubation temperatures on post-hatching survival, growth, and critical thermal maximum of larval cisco (*Coregonus artedi*) sampled from lakes Superior and Ontario. Embryos were incubated at water temperatures of 2.0, 4.4, 6.9, and 8.9°C to simulate present and increased winter temperatures, and hatched larvae were reared in a common environment. For both populations, larval survival and critical thermal maximum were negatively related to incubation temperature, and larval growth was positively related to incubation temperature. The magnitude of change across incubation temperatures was greater in the population sampled from Lake Superior than Lake Ontario for all traits examined. The more rapid decrease in survival and critical thermal maximum across incubation temperatures for larval cisco in Lake Superior, compared to those from Lake Ontario, suggests that Lake Superior larvae may possess a more limited ability to acclimate to and cope with increasing winter water temperatures. However, the rapid increase in growth rates across incubation temperatures in Lake Superior larvae suggests they could recover better from hatching at a small length induced by warm winters, as compared to Lake Ontario larvae. Our results suggest propagation and restoration programs may want to consider integrating natural habitat preferences and maximizing phenotypic variability to ensure offspring are set up for success upon stocking.

## Introduction

Water temperatures are rising around the globe (Austin and Colman, 2008; Maberly et al., 2020; O’Reilly et al., 2015; Woolway et al., 2020) and poses a threat to ectotherms, such as fish, that have limited thermal tolerance ranges (Comte and Olden, 2017; Dahlke et al., 2020; Little et al., 2020). Thermal tolerances vary with ontogenetic development (Dahlke et al., 2020; Sunday, 2020) and affect reproduction, metabolic rates, growth, and overall survival (Brett, 1979; Brown et al., 2004; Busch et al., 2012; Gillooly et al., 2002; Little et al., 2020; Ohlberger et al., 2007). Vulnerability of fishes to climatic warming is highest for cold stenothermic species that lack the ability to migrate to suitable temperatures. Specific vulnerability of local populations will likely depend on future climate regime shifts and the temperature requirements of spawners and embryos (Dahlke et al., 2020; Sunday, 2020). In the short-term, lacustrine spawners may cope with warming waters by shifting spawning timing or using deeper and colder spawning habitat, or potentially in the long-term through thermal adaptation across generations. Adaptation, however, may be too slow to keep pace with changing thermal conditions (Bruge et al., 2016). For autumn spawners, spawning later in the season after waters have cooled sufficiently may still impact embryo development due to warmer winter temperatures and earlier spring warming.

Freshwater whitefishes, Salmonidae Coregoninae (hereafter coregonines), are cold, stenothermic fishes of ecological and socio-economic importance throughout the northern hemisphere (Elliott and Bell, 2011; Isaak, 2014; Jeppesen et al., 2012; Jonsson and Jonsson, 2014; Karjalainen et al., 2015; Stockwell et al., 2009). In the Laurentian Great Lakes, cisco (*Coregonus artedi*) was historically the most abundant ciscoe (*sensu* Eshenroder et al., 2016) species, a primary prey fish of lake trout (*Salvelinus namaycush*), and a commercial fishing target since the early 1800s (Bogue, 2001; Chiarappa, 2005). Most cisco spawning stocks collapsed by the mid-1900s (Baldwin et al., 2009; Koelz, 1929). Lake Superior stocks partially recovered by the early-1990s (Stockwell et al., 2009), but contemporary abundance is considered to be below historical levels (Rook et al., 2021). Present Lake Superior cisco population abundance is hypothesized to be limited by reduced and inconsistent survival of fish to age-1 due to climatic warming over the past two decades (Van Cleave et al., 2014) and lower overall ecosystem productivity due to reduced phosphorus inputs as compared to 1900-1970 (Rook et al., 2021). Variable and weak year-class strength of coregonines has been observed worldwide over the past several decades and has been associated with annual variations in lake ice formation and winter-spring thermal conditions (Anneville et al., 2015; Karjalainen et al., 2015; Marjomäki et al., 2004; Nyberg et al., 2001).

Most coregonines spawn nearshore in late-autumn, embryos incubate under ice, and hatch in spring near ice-out, when rising spring water temperatures trigger hatching (Karjalainen et al., 2021, 2019, 2015; Stockwell et al., 2009). Increases in air temperature and water temperatures of seasonally ice-covered lakes are projected to be greatest during the winter and spring, respectively, in response to climate change (Christensen et al., 2007; Ozersky et al., 2021; Schindler et al., 1990; Winslow et al., 2017).

The larval period of fishes is critical for year-class success (Cushing, 1990; Hjort, 1914), but the physiological effects of thermal stress from non-optimal embryo incubation temperatures on post-hatching survival are unclear. Additional physiological pressures as a result of warming winter temperatures could be detrimental. The match-mismatch hypothesis postulates that larval survival is dependent on a temporal and spatial match between larval feeding capabilities, such as swimming ability and prey acquisition, and prey availability (Cushing, 1990). Warmer incubation temperatures lead to earlier hatch dates and altered morphological developments, such as smaller lengths and larger yolk sacs, that reduce larval feeding efficiency (Darowski et al., 1988), compared to colder incubated embryos (Karjalainen et al., 2015; Stewart et al., 2021a). The selective pressures from elevated temperatures on embryonic and larval coregonine development and survival may lead to adaptation, but the thermal trigger for the response and the mechanism of the response are unknown. Consequently, quantifying the potential response and adaptive capacity of cisco to warming winter and spring water temperatures is needed.

We experimentally evaluated how cisco embryo incubation temperatures influenced the survival and performance of hatching larvae within and between two Great Lakes cisco populations. We hypothesized that warmer, sub-optimal cisco embryo incubation temperatures decrease larval survival, growth, and critical thermal limits compared to embryo incubation temperatures that mimic cold, pre-climate change conditions. If our hypothesis is supported, we would expect a negative relationship between embryo incubation temperature and the larval traits examined for wild cisco populations when reared artificially.

## Methods

### Ethics

All work described here was approved for ethical animal care under University of Vermont’s Institutional Animal Care and Use Committee (Protocol # PROTO202000021).

### Crossing Design and Fertilization

Cisco were collected from the Apostle Islands, Lake Superior (46.85°, −90.55°) and Chaumont Bay, Lake Ontario (44.05°, −76.20°) in December 2019. Eggs and milt were stripped from 12 females and 16 males from each population and artificially fertilized under a blocked, nested full-sib, half-sib fertilization design to create a maximum of 48 families. A single fertilization block consisted of four males each paired to three unrelated females, where all offspring of a given female were full siblings (Stewart et al., 2021a).

For clarity, our operational use of a population represents a single species sampled from a single location within a single lake (*e.g.,* cisco from the Apostle Islands region in Lake Superior). Our sampling efforts represent a single location within large lakes and does not likely capture the possible genetic variation within a species or population.

### Rearing Conditions

Full embryo incubation methods are described in Stewart et al. (2021). Embryos were incubated in 24-well cell culture microplates placed in climate-controlled chambers (Memmert^®^ IPP260Plus) at mean (SD) constant temperatures of 2.0 (0.5), 4.4 (0.2), 6.9 (0.2), and 8.9 (0.3)°C. These temperatures were selected to mimic present and potentially warmer winter temperatures (Titze and Austin, 2014) at typical cisco spawning depths (<100 m, Goodyear, 1982). Reconstituted freshwater medium was used during fertilizations and incubations (International Organization For Standardization 6341, 2012) to standardize the chemical properties of the water among all treatments and between populations. After hatching, larvae were photographed alive and ventrally (Nikon^®^ D5600 and Nikon^®^ AF-S DX 18-55mm lens). Total length was measured from images using Olympus^®^ LCmicro.

Newly-hatched larvae were transferred to rearing tanks segregated by population and incubation temperature. Larvae from Lake Superior were reared in four (4 incubation treatments × 1 replicate) 150-liter oval tanks. Larvae from Lake Ontario were reared in eight (4 incubation treatments × 2 replicates) 150-liter oval tanks. Lake Ontario larvae were divided equally by families (*i.e.,* up to 24 of 48 total larvae per family per replicate tank) into replicate tanks per incubation temperature treatment. Lake Superior larvae were unreplicated as a result of low fertilization success and embryo survival - insufficient numbers were available for multiple rearing tanks. All rearing tanks were supplied with chilled, recirculating water maintained at 6.5°C (mean (SD) = 6.36 (1.17)). Water temperatures (±0.2°C) were recorded hourly. Larvae in all rearing tanks were exposed to the same photoperiod cycle (*i.e.,*12-hr light, 12-hr dark) with gradual sunrise and sunset transitions. Full spectrum (*i.e.,* 380-780 nm), white LED lights (AquaShift^®^ MLA-WH) were used to simulate daylight. Dead larvae were removed and counted each day. Larvae were fed *Artemia* and transitioned to Otohime A dry feed one-week post-hatch. Food was provided *ad libitum*.

### Thermal Challenge

After 60 days, larvae from each population, incubation treatment, and replicate rearing tank were thermally challenged. Because larvae within and among rearing tanks did not hatch on the same day, 60-days post-hatch was calculated from the date of 50% hatching for each rearing tank. Larvae were transferred to 5.4-liter clear, rectangular tanks, with two replicate tanks per rearing tank and approximately 50 larvae, or as many available, were used in each replicate tank. Water temperature was 10°C and larvae were allowed to acclimate to this temperature for 12 hours prior to the thermal challenge. The water in the thermal challenge system was recirculated among all replicate tanks and aerated. During the thermal challenge, water temperatures were raised from 10.0°C at a constant rate of 0.5°C per 30 minutes until all larvae were deceased. Larvae were considered terminated when loss of equilibrium was achieved and were motionless for at least 5 seconds. Once endpoint criteria were met, larvae were euthanized, photographed, and preserved in 95% ethanol. The elapsed time and temperature at termination of each individual larvae was recorded and total length was measured from the images.

All larvae from the 8.9°C treatment died during the acclimation period from an unknown cause, thus, only thermal challenge data from 2.0, 4.4, and 6.9°C are presented.

### Statistical Analyses

All statistical analyses were performed in R version 4.0.5 (R Core Team, 2021).

Larval survival was estimated for each rearing tank, across all families, as the percent of larvae surviving between hatching and 60 days after the date of 50% hatching. Our estimates of larval survival from Lake Superior are unreplicated. However, useful information can still be gleaned without strict statistical testing (*e.g.*, Davies and Gray, 2015). Observations of single estimates of larval survival across incubation temperatures could foster further hypotheses and lead to more focused studies.

Similar to larval survival estimates, larval growth rate estimates for Lake Superior were unreplicated. To this end, we qualitatively compared absolute growth rates between populations and across incubation temperatures by generating bootstrapped confidence intervals for the observed absolute growth rate estimates. For each population, incubation temperature treatment, and replicate rearing tank, a bootstrapped mean length-at-hatch was calculated from random sampling with replacement from the observed lengths-at-hatch, and a bootstrapped mean final length was calculated from random sampling with replacement from the observed final lengths. The difference between the bootstrapped mean final length and bootstrapped mean length-at-hatch was calculated and divided by the duration of the larval experiment (*i.e.,* absolute growth rate). The bootstrap procedure was repeated 10,000 times. The bootstrapped absolute growth rate distributions were used to calculate the 2.5 and 97.5 percentile values (*i.e.,* 95% confidence interval) as a measure of variation around the observed absolute growth rate, and to qualitatively assess the likelihood of differences in growth among populations and incubation temperature treatments, in absence of replication. For Lake Ontario, the 95% confidence intervals were calculated as the mean 2.5 and 97.5% percentiles across replicate tanks. Comparisons were made by examining the overlap of the observed mean absolute growth rate to the bootstrapped 95% confidence intervals of all pairwise comparisons.

The critical thermal maxima (CTMax) of larval cisco from each population and incubation temperature treatment was expressed as the arithmetic mean of the temperature at which endpoint criteria were reached (Mora and Ospina, 2001). Although we have estimates for each individual larva within replicate thermal challenge tanks, the larvae from Lake Superior were reared in a single rearing tank and thus the estimates are not independent and cannot be treated as true replicates. Therefore, a similar bootstrap approach as described for larval growth was used to qualitatively compare CTMax among populations and incubation temperature treatments. For each population, incubation temperature treatment, and replicate rearing tank, we generated a bootstrap sample by randomly selecting, with replacement, a termination temperature n times, where n equals the number of observations in the experiment. The CTMax was calculated for each bootstrapped sample and the distribution of bootstrapped CTMax was used to calculate the 95% confidence interval as a measure of variation around the observed CTMax. The bootstrap procedure was repeated 10,000 times. For Lake Ontario, the 95% confidence intervals were calculated as the mean 2.5 and 97.5% percentiles across replicate tanks. Comparisons were made by examining the overlap of the observed mean CTMax to the bootstrapped 95% confidence intervals of all pairwise comparisons.

## Results

### Larval Survival

A total of 9,605 larvae hatched and were reared from lakes Superior (2,332 larvae) and Ontario (7,273 larvae) across all incubation temperatures. Larval survival was highest at the 2.0°C incubation temperature and decreased with warming incubation temperatures for both populations (Figure 1). Survival rates were 38.7% at 2.0°C, 17.7% at 4.4°C, 1.1% at 6.9°C, and 5.4% at 8.9°C for Lake Superior larvae and 43.3% at 2.0°C, 35.3% at 4.4°C, 12.4% at 6.9°C, and 2.6% at 8.9°C for Lake Ontario larvae. Larval survival was higher for Lake Ontario larvae than Lake Superior larvae across all incubation temperature treatments, except 8.9°C. Lake Ontario larvae had similar survival rates (< 9% difference) at the 2.0 and 4.4°C incubation temperatures, whereas Lake Superior larval survival decreased 21% from the 2.0° to 4.4°C incubation temperatures (Figure 1).

**Figure 1.**
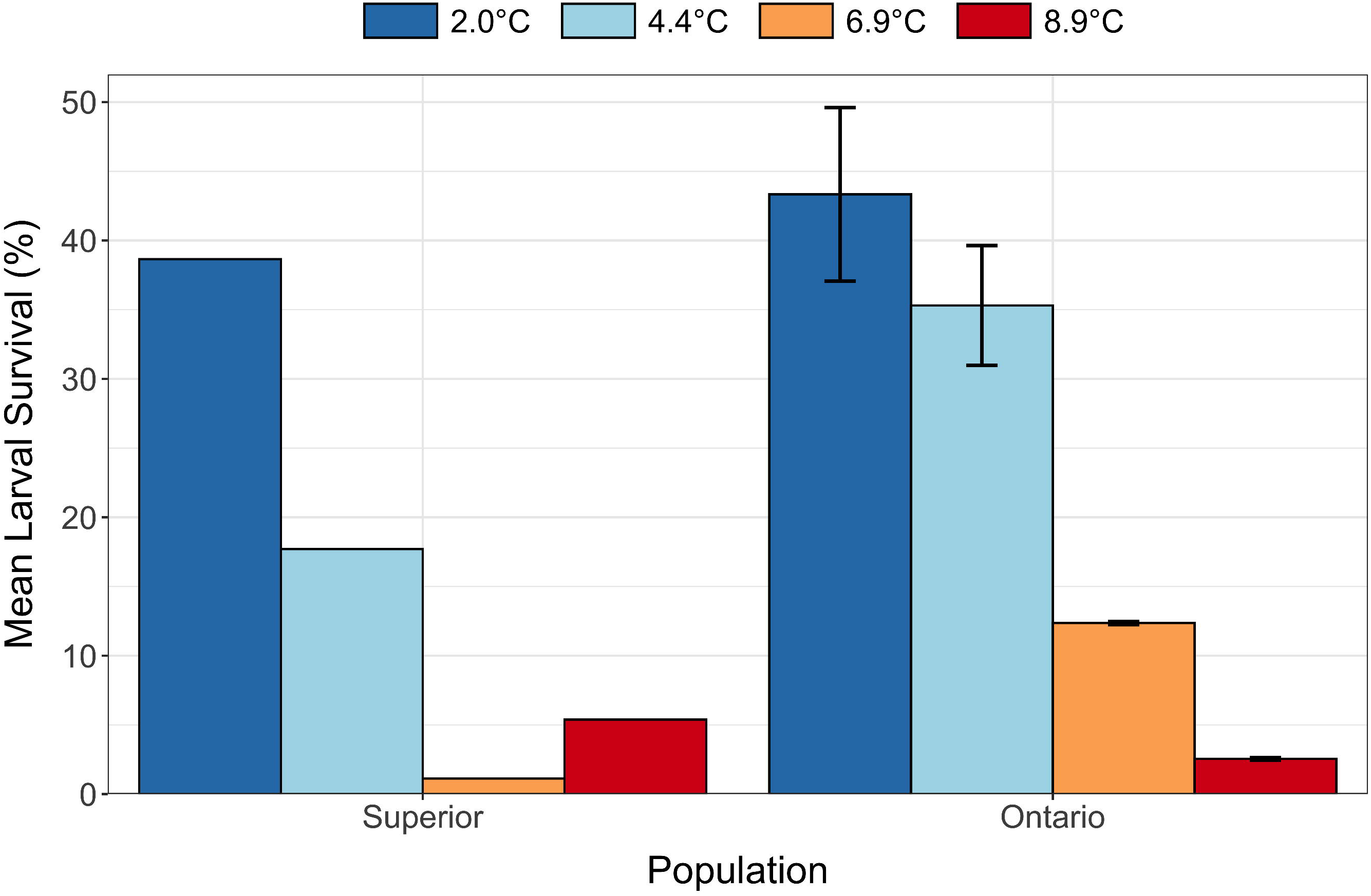
Mean larval survival (%) for larval cisco (*Coregonus artedi*) from Lakes Superior and Ontario incubated at 2.0, 4.4, 6.9, and 8.9°C across replicate rearing tanks. Error bars indicate standard error. Lake Superior mean survival estimates are unreplicated and thus do not have error estimates.

### Larval Growth

Larval cisco absolute growth rates increased with warming incubation temperatures in both populations (Figure 2). Larvae from Lake Superior had lower absolute growth rates at 2.0 and 4.4°C (0.049 and 0.044 mm day^−1^, respectively) compared to Lake Ontario (0.056 and 0.061 mm day^−1^, respectively). Absolute growth rates increased at 6.9°C for Lake Superior (0.057 mm day^−1^) and 8.9°C for Lake Ontario (0.078 mm day^−1^), and both populations had similar absolute growth rates at 6.9 and 8.9°C (mean difference <0.001 and 0.012 mm day^−1^, respectively; Figure 2).

**Figure 2.**
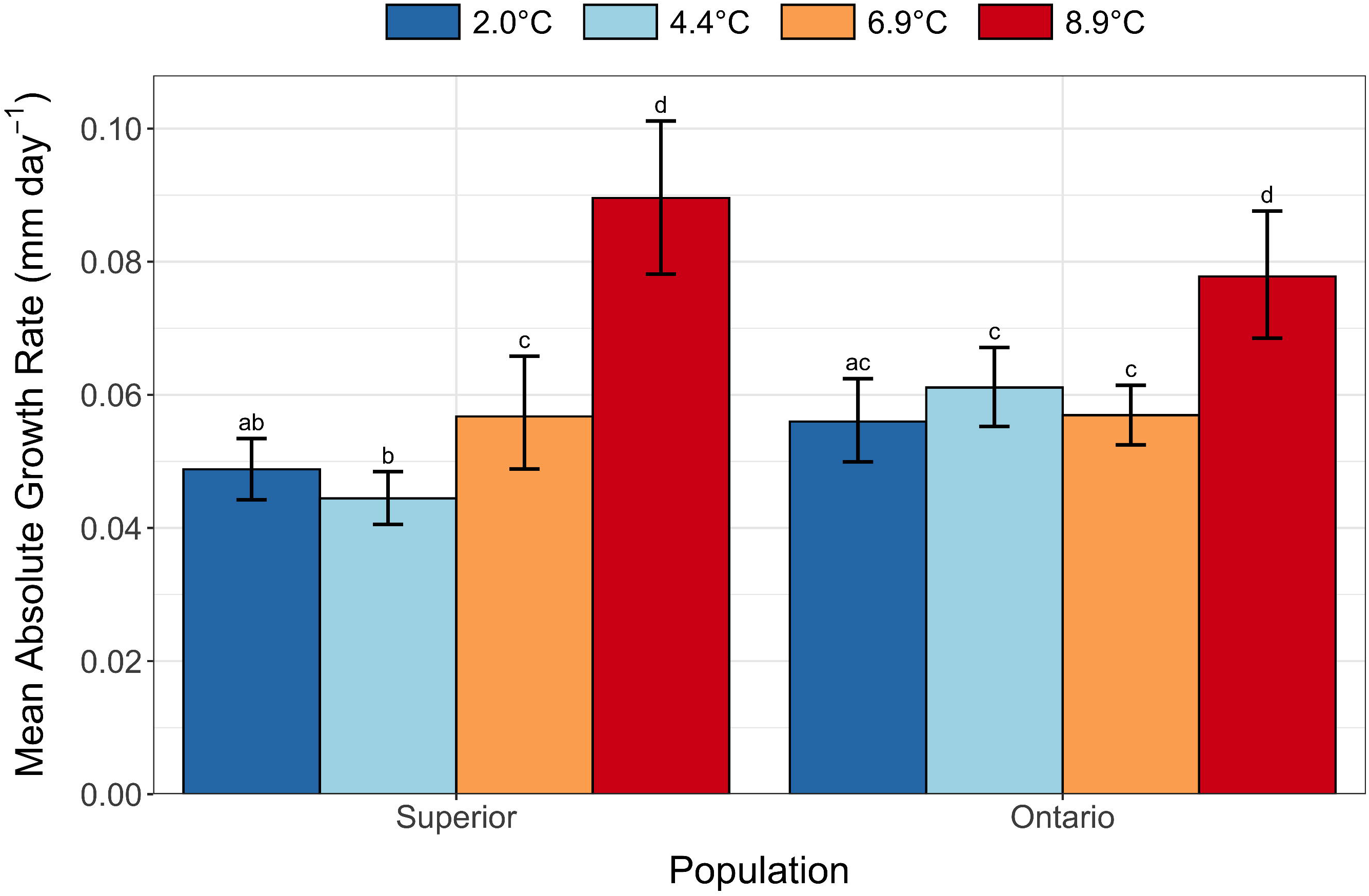
Mean absolute growth rates (mm day^−1^) for larval cisco (*Coregonus artedi*) from lakes Superior and Ontario incubated at 2.0, 4.4, 6.9, and 8.9°C. Error bars indicate 95% bootstrapped confidence intervals. Letters indicate overlap of the observed mean absolute growth rate to the bootstrapped 95% confidence intervals of all pairwise comparisons.

### Thermal Challenge

Critical thermal limit in larval cisco decreased with warming incubation temperatures in Lake Superior and Lake Ontario (Figure 3). Larvae from Lake Superior incubated at 2.0°C had the highest CTMax (25.81°C). However, CTMax in Lake Superior decreased by 0.83 and 0.77°C between the 2.0 to 4.4°C and the 4.4 and 6.9°C incubation temperature treatments, respectively. CTMax was similar for Lake Ontario larvae incubated at 2.0 and 4.4°C (24.99 and 24.96°C, respectively) and decreased at 6.9°C (24.67°C).

**Figure 3.**
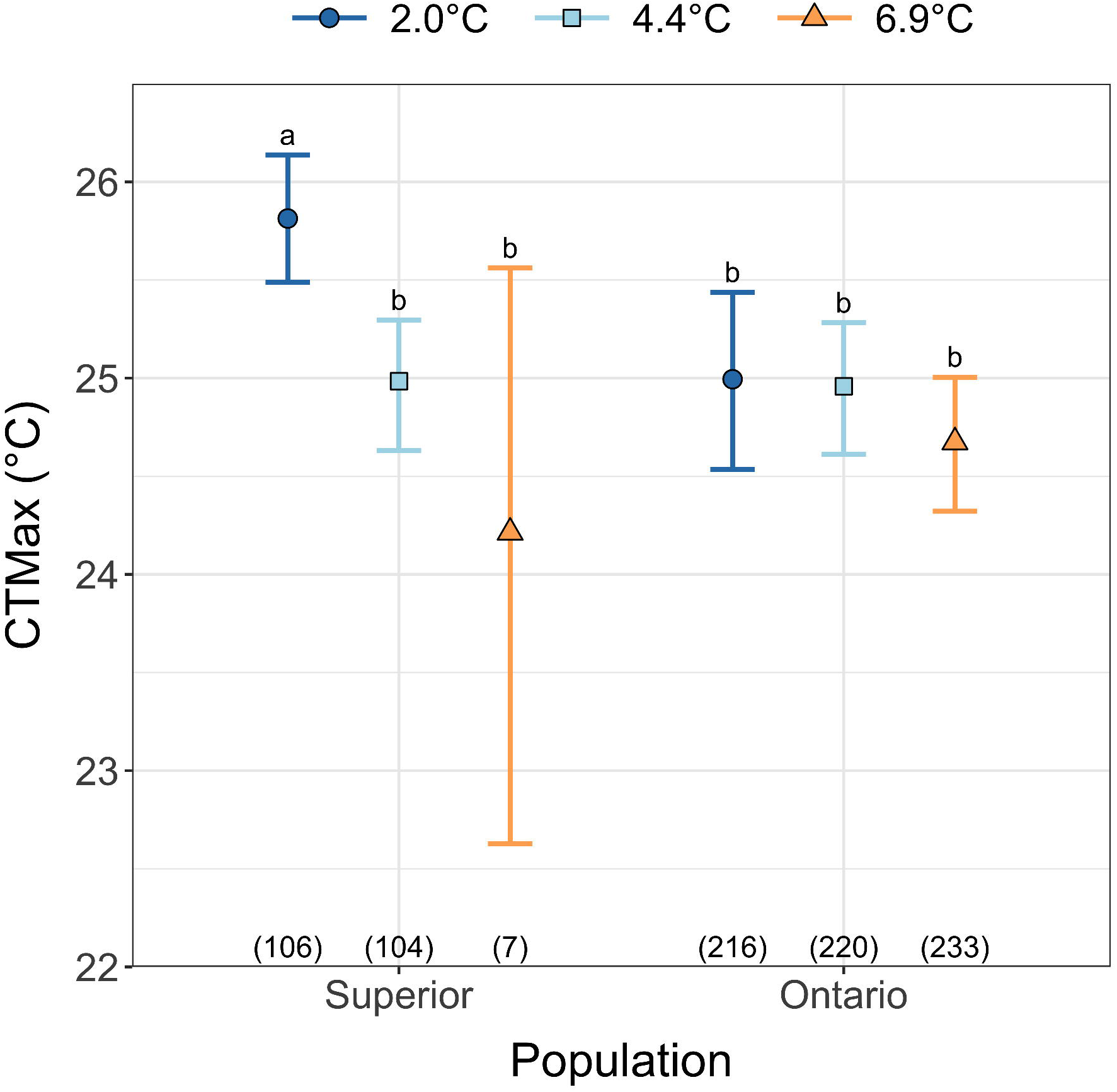
Critical thermal maxima (CTMax; °C) for larval cisco (*Coregonus artedi*) from lakes Superior and Ontario incubated at 2.0, 4.4, and 6.9°C. Error bars indicate 95% bootstrapped confidence intervals. Letters indicate overlap of the observed CTMax to the bootstrapped 95% confidence intervals of all pairwise comparisons. Sample sizes are indicated in parentheses.

## Discussion

Survival, growth rates, and critical thermal limits of cisco larvae from lakes Superior and Ontario were influenced by embryo incubation temperatures that were warmer than current natural winter water temperatures in these lakes. First, we found a negative relationship between larval survival and embryo incubation temperature. Second, warmer embryo incubation temperatures increased larval growth rates. Third, critical thermal limits decreased with warming incubation temperatures. Lastly, the magnitude of change across incubation temperature treatments was greater in cisco from the historically colder Lake Superior than Lake Ontario for all traits examined. These results suggest increased risk to Laurentian Great Lakes cisco populations in response to projected climatic warming.

Our hypothesis that larval survival is highest at the coldest incubation temperature, which mimicked the natural lake environment, was supported. Between the two lakes, Lake Superior cisco had a sharper decline in larval survival from 2.0 to 4.4°C compared to Lake Ontario cisco. Though both populations are cold adapted, the result suggests Lake Superior cisco were more cold-water adapted than those from Lake Ontario. Lake Superior is colder than Lake Ontario (Mason et al., 2016; Millar, 1952) and spawning cisco from Lake Superior were collected at an open lake location, whereas spawning cisco from Lake Ontario cisco were collected in a shallow, protected bay. Water temperatures in shallower protected habitats increase more rapidly after ice-out and have higher maximum spring and summer temperatures (*i.e.,* Lake Ontario sampling location; Minns et al., 2011) compared to deeper, open-water locations (*i.e.,* Lake Superior sampling location; Titze and Austin, 2014) because the heat capacity of water is positively related to depth and water is mixed less in protected bays (Assel et al., 2003; Gan and Liu, 2020; Verburg and Antenucci, 2010). Interactions among winter and spring temperatures, hatching dates, zooplankton availability and larval size-dependent predation mortality influence year-class strength of vendace (*C. albula*) and European whitefish (*C. lavaretus*) in Europe (Anneville et al., 2009; Marjomäki et al., 2004; Mehner et al., 2011; Miller et al., 1988). Spring warming rates in particular appear to play a critical role in prey availability and larval growth and survival of autumn-spawning coregonines (Karjalainen et al., 2015; Myers et al., 2014).

The transition from endogenous to exogenous feeding is critical to larval fish survival (Cushing, 1990; Hjort, 1914). Higher winter temperatures induce earlier coregonine embryo hatching and cause larvae to have smaller lengths-at-hatch and larger yolk-sac volumes (Karjalainen et al., 2015; Stewart et al., 2021a). Larvae hatching with larger yolk sacs may have more time to transition to exogenous feeding (Hjort, 1914; Lucke et al., 2020; Miller et al., 1988), but at a cost to swimming efficiency and predator avoidance (Darowski et al., 1988; Myers et al., 2014). In wild populations, earlier hatching may also increase the mismatch between the onset of spring plankton blooms and larval prey, increasing the risk for starvation and higher larval mortality (Cushing, 1990; Myers et al., 2014). Interactions among winter and spring temperatures, hatching dates, zooplankton availability and larval size-dependent predation mortality influence year-class strength of vendace (*C. albula*) and European whitefish (*C. lavaretus*) in Europe (Anneville et al., 2009; Marjomäki et al., 2004; Mehner et al., 2011; Miller et al., 1988). Spring warming rates in particular appear to play a critical role in prey availability and larval growth and survival of autumn-spawning coregonines (Karjalainen et al., 2015; Myers et al., 2014). Our experiment provided cisco larvae a predator-free environment with *ad libitum* food immediately after hatching, yet we still observed sharp declines in larval survival for those incubated at increased temperatures. We suggest an additional or alternative hypothesis for a survival bottleneck under climate change scenarios is that larval cisco survival may not be as limited by prey availability but instead by reduced physiological condition caused by warmer embryo incubations.

Rapid larval growth is associated with high survival (Blaxter, 1986; Houde, 1989; Miller et al., 1988; Myers et al., 2014; Ware, 1975). In our experiment, larval cisco exhibited low survival despite higher absolute growth rates when incubated at warmer temperatures. These results did not support our hypothesis that warmer, sub-optimal cisco incubation temperatures decrease larval growth rates. Coregonine embryos incubated at high temperatures (i.e., > 6°C) often hatch prematurely and are underdeveloped (Colby and Brooke, 1970; Price, 1940). Warmer incubations may require free-floating embryos to rapidly convert yolk for development. In this sense, higher absolute growth rates gained from warming incubation temperatures may not be optimal for survival. Larval cisco from Lake Superior use a mixed-feeding strategy with endogenous energy reserves (i.e., yolk) and exogenous feeding overlapping at lengths between 10.0-12.0 mm (Lucke et al., 2020). Embryos incubated at colder temperatures (i.e., 2.0 and 4.4°C) from the sampled populations of lakes Superior and Ontario cisco had mean lengths-at-hatch from 9.9-11.3 mm, whereas mean lengths-at-hatch ranged from 8.7-9.7 mm at the warmest incubation temperature (8.9°C; Stewart et al., 2021). A combination of field and experimental data suggests that cold, long incubations with prolonged development results in larger length-at-hatch with less endogenous energy reserves, the ability to immediately use a mixed endogenous and exogenous feeding strategy, and lower growth rates could be the best scenario to maximize larval survival (Lucke et al., 2020; Stewart et al., 2021a). However, this ‘goldilocks scenario’ may only work if all biotic and abiotic conditions (*e.g.,* water temperature, appropriately sized prey, etc.) match cisco phenotypes. This hypothesis remains to be tested, as our experiments provided a stable and optimal temperature and feeding environment, conditions that cannot be assumed to occur in the wild.

The ability of larval cisco to use favorable nursery habitat near the lake surface is directly related to their ability to tolerate spring-summer surface water temperatures. The increase in CTMax with decreased incubation temperature supported our hypothesis that cold, pre-climate change conditions would maximize thermal performance. The different magnitudes of change between cisco from lakes Superior and Ontario could be explained from evolutionary adaptations to local conditions. Fish populations from high-latitude, low-temperature locales often compensate for slower metabolism and lower growth rates by having more efficient physiological performance than low-latitude populations (*i.e.,* countergradient variation; (Conover and Present, 1990; Jonassen, 2000; Reist et al., 2006). Lake Superior experiences colder and less seasonal variation in water temperature than Lake Ontario (Calamita et al., 2021; Zhang et al., 2018), and larval cisco from Lake Superior may have more efficient physiological adaptations (*e.g.,* cardiac and respiratory performance) which could explain the high thermal tolerance at cold incubation temperatures and sensitivity to increased temperatures. Our results suggest research on mechanisms driving the observed differences in CTMax between populations (*e.g.,* cardiac failure, oxidative damage to tissue, body mass, stress biomarkers, protein denaturation, etc.) may prove insightful.

Our results have implications for current and proposed hatchery-based restoration efforts of coregonines in the Laurentian Great Lakes (Bronte et al., 2017). We found that cisco offspring from two of the Great Lakes raised at warm incubation temperatures (*i.e.,* > 4.5°C) had lower overall performance than individuals incubated at cold temperatures (*i.e.,* < 4.5°C). Many coregonine hatchery facilities around the Great Lakes do not or cannot incubate embryos under natural lake thermal conditions (*i.e.,* cold water temperatures, < 4.5°C; Bronte et al., 2017). Hatchery-produced fish can have lower fitness in natural environments than wild fish (Araki et al., 2008; Bailey et al., 2010; Christie et al., 2014). Offspring from parents haphazardly selected for artificial breeding and reared in captivity before release have the potential to induce strong directional selection and harm naturally recruiting populations (Araki and Schmid, 2010; Tingley III et al., 2019). Transgenerational effect of lower larval performance and its potential effect on the response to selection are unknown but warrants investigation (Araki et al., 2008; Araki and Schmid, 2010; Christie et al., 2014; Ford, 2002). The consequences an artificial environment may have on the genetic diversity within a population and fitness of post-stocking individuals needs to be considered in ongoing restoration and conservation efforts (Tingley III et al., 2019).

Identifying the genetic mechanisms (*i.e.,* SNPs and gene expression) involved in the thermal adaptation and acclimation of coregonine populations is an important next step. Variation in certain genetic markers and survival under thermal stress may allow managers to determine the genotypes associated with increased survival at variable or increasing temperatures (Narum et al., 2013). Examining gene expression across populations and temperature treatments will help identify and evaluate the function of differentially expressed genes and potential physiological pathways that may be disproportionately under- or over-represented with thermal stress (Rougeux et al., 2018). Furthermore, the combination of genomic tools (*e.g.,* genome-wide association study and RNA-seq) in thermal ecology experiments can provide valuable insight into the functional significance of markers associated with thermal tolerance (Rougeux et al., 2018). Considerable progress has been recently made in advancing our genomic knowledge of Laurentian Great Lakes coregonines that will provide a foundation for this work (Ackiss et al., 2020; Blumstein et al., 2020; Eaton et al., 2021; Lachance et al., 2021).

## Conclusion

The rapidity at which winter environments are changing has revealed our ‘blind spot’ for winter biology (Ozersky et al., 2021). The results presented here and elsewhere (Karjalainen et al., 2016, 2015; Stewart et al., 2021a, 2021b) focus on how coregonine reproduction may be impacted by a warming climate and suggest that while we have much to learn, the effects of warming winters will vary among populations and with the magnitude of warming. These results highlight the importance of integrating natural habitat preferences into stock propagation programs to ensure offspring are set up for success upon reintroduction. A challenge for managers and propagation facilities is to consider the impact embryo incubation conditions may have on larval survival and performance in relation to production targets. Additionally, propagation and stocking may accomplish the short-term restoration objective of supplementing wild populations, but other limiting factors (*e.g.,* habitat loss, anthropogenic disturbances, water quality, invasive species) also need to be addressed to achieve long-term population conservation and viability (Tingley III et al., 2019). Maximizing phenotypic variation and adaptability to changing conditions (*i.e.,* portfolio effect; Schindler et al., 2015, 2010) is a strong consideration in restoration and conservation efforts. Embracing management strategies that foster increased early-life stage fitness could improve the ability of coregonines to cope with environmental change in the wild and aid in addressing recruitment bottlenecks.

## Acknowledgments

We thank staff at the Wisconsin Department of Natural Resources Bayfield Fisheries Field Station, U. S. Geological Survey (USGS) Tunison Laboratory of Aquatic Science, and New York State Department of Environmental Conservation Cape Vincent Fisheries Station for field collections of spawning adults. Rachel Taylor, Dan Yule, and Caroline Rosinski helped with fertilizations and incubation experiment maintenance. [insert name] provided the USGS solicited review that strengthened the manuscript, as did anonymous peer reviewers and Stockwell and Dr. Ellen Marsden laboratory members. This work was funded by the USGS [grant/cooperative agreement number G16AP00087 and G17AC00042] to the Vermont Water Resources and Lakes Studies Center and the University of Vermont. Additionally, this work was made possible with funds made available to Lake Champlain by Senator Patrick Leahy through the Great Lakes Fishery Commission. Any use of trade, product, or firm names is for descriptive purposes only and does not imply endorsement by the U.S. Government.

